# The Toll-Like Receptor 5 agonist flagellin prevents *Non-typeable Haemophilus influenzae*-induced exacerbations in cigarette smoke-exposed mice

**DOI:** 10.1101/2020.07.08.193128

**Authors:** Magdiel Pérez-Cruz, Bachirou Koné, Rémi Porte, Christophe Carnoy, Julien Tabareau, Pierre Gosset, François Trottein, Jean-Claude Sirard, Muriel Pichavant, Philippe Gosset

## Abstract

Chronic obstructive pulmonary disease (COPD) is a major cause of morbidity and mortality worldwide. The major bacterial cause of COPD exacerbations is non-typeable *Haemophilus influenzae* (NTHi). This susceptibility to infection involves a defective production of interleukin (IL)-22 which plays an important role in mucosal defense. Prophylactic administration of flagellin, a Toll-like receptor 5 (TLR5) agonist, protects healthy mice against respiratory pathogenic bacteria. We hypothesized that TLR5-mediated stimulation of lung immunity might prevent COPD exacerbations due to NTHi. Mice were chronically exposed to cigarette smoke and then infected with NTHi. According our preventive or therapeutic protocol, flagellin was administered intraperitoneally. Cigarette smoke-exposed mice treated with flagellin showed a lower bacterial load in the airways, the lungs and the blood. This protection was associated with an early neutrophilia, a lower production of pro-inflammatory cytokines and an increased IL-22 production. Flagellin treatment decreased the recruitment of inflammatory cells and the lung damages related to exacerbation. Protective effect of flagellin against NTHi was altered by treatment with anti-IL-22 blocking antibodies in cigarette smoke-exposed mice and in *Il22*^*−/−*^ mice. Flagellin treatment also amplified the production of the β-defensin2 anti-bacterial peptides. This study shows that stimulation of innate immunity by a TLR5 ligand is a potent antibacterial treatment in cigarette smoke exposed mice, suggesting innovative therapeutic strategies against acute exacerbation in COPD.

## Introduction

Chronic obstructive pulmonary disease (COPD) is characterized by a progressive and irreversible decline in lung function ^1^. Being the third leading cause of death worldwide, it is mainly caused by chronic exposure to cigarette smoke (CS) or pollutants ^2^. Inhalation of CS essentially leads to activation of epithelial cells and macrophages responsible for the mobilization of effector and immuno-modulatory cells including neutrophils and natural killer T (NKT) cells ^3, 4^. The chronic inflammatory response progressively leads to airway remodeling, impaired mucociliary clearance and parenchymal destruction in the lungs, further culminating in irreversible airflow limitation ^5^. These components are involved in the increased susceptibility of COPD patients to bacterial and viral airway infections.

Airway colonization with bacteria such as *Haemophilus influenzae, Streptococcus pneumoniae* and *Moraxella catarrhalis* contributes to the pathogenesis and clinical course of the disease ^6^. This colonization is responsible for lung infection leading to exacerbations of the disease, which have a strong impact on health status, exercise capacity, lung function, and mortality. Non-typeable *Haemophilus influenzae* (NTHi), a Gram-negative coccobacillus that lacks a polysaccharide capsule, is an important cause of COPD exacerbations and comorbidity ^7, 8^. Acute exacerbations invariably scarred the chronic course of COPD ^9^. Bacterial infections are first controlled by the innate immune system, which implicated pathogen-associated molecular pattern (PAMP) recognition by Toll-like receptors (TLR) such as those recognizing flagellin (TLR5) responsible for the mobilization of effector cells ^10^. During COPD, bacterial infection is characterized by an increased influx of immune cells, including neutrophils, macrophages, dendritic cells (DC) and T lymphocytes ^3, 11, 12^. However, this response is not effective enough to clear the pathogens. In this context, we recently reported a defective production of IL-22 in response to bacteria both in COPD patients and mice chronically exposed to CS, whereas IL-17 production is only altered after infection by *S. pneumoniae* ^13, 14^. Interestingly, the Th17 cytokines IL-17 and IL-22 promote the recruitment of neutrophils, the synthesis of antimicrobial peptides and the expression of tight junction molecules ^15, 16^, a mechanism explaining the essential role of IL-22 in the clearance of NTHi ^17^. Supplementation of COPD mice with recombinant IL-22 increases the clearance of the bacteria and prevents the development of COPD exacerbations in mice. Several reports showed that activation of innate receptors, including TLR, is able to elicit protective immune responses against infections ^18, 19^. Among them, systemic administration of flagellin, the main component of bacterial flagella and the TLR5 ligand, induces immediate production of Th17 cytokines through the activation of DC and type 3 innate lymphoid cells ^20^.

In this study, we hypothesized that systemic administration of flagellin could limit the development of NTHi-induced COPD through eliciting an appropriate protective IL-22 response. We reported here that systemic stimulation of the innate immunity by flagellin from *Salmonella enterica* serovar Typhimurium (FliC) could prevent COPD exacerbation induced by NTHi. We also showed that the protective effect of flagellin against NTHi was dependent of IL-22 and associated with the upregulation of anti-microbial peptides.

## Material and Methods

### Animals

Male C57BL/6 (WT) mice, 6–8 weeks old were purchased from Janvier Labs (Le Genest-St-Isle, France). *Il22*^*−/−*^ mice were obtained from Jean-Christophe Renauld (Brussel, Belgium). WT mice were daily exposed to cigarette smoke (CS) during 12 weeks (5 cigarettes/day, 5 days / week) to mimic COPD pathogenesis ^4^. 3R4F research cigarettes were obtained from the University of Kentucky Tobacco and Health Research Institute (Lexington, KY, USA). The control group was exposed to ambient air. After 12 weeks of CS or air exposure, mice were either treated intranasally with phosphate buffered saline (PBS) or NTHi (n=4 per group). *Il22*^*−/−*^ mice were infected or not with NTHi. Experiments were performed at least in triplicate. Mice were examined every working day and humane endpoints essentially based on an important weight loss of more than 20% were applied for each experiment. However, our protocols did not provoke such a situation. Mice were euthanized by cervical dislocation according to the French government guidelines of laboratory animal care and approved by the Departmental Direction of Veterinary Services (Prefecture of Lille, France), European guidelines of laboratory animal care (number 86/609/CEE) and French legislation (Government Act 87-848). The present project has been approved by the national Institutional Animal Care and Use Committee (CEEA 75) and received the authorization number APAFIS# 7281.

### Mice infection and flagellin treatment

NTHi 3224A strain was grown to log-phase in brain-heart infusion (BHI) broth (AES Laboratory) supplemented with 10μg/ml haematin and 10μg/ml nicotinamide adenine dinucleotide (NAD) (SIGMA, St Louis, MI, USA), and stored à −80°C in BHI 10% glycerol for up to 3 months.

Working stocks were thawed, washed with sterile PBS, and diluted to the appropriate concentration. Mice were anesthetized and intranasally (i.n.) infected with 2.5×10^6^ CFU of NTHi. The number of infectant bacteria was confirmed by plating serial dilutions onto chocolate agar plates.

To prepare heat-killed (HK) NTHi, bacteria were grown to a log-phase (O.D_600nm_=0.7−0.8 units) and inactivated for 1 hour at 56° C in a hot-water-bath. Broth cultures were then plated onto chocolate agar plates and incubated overnight to check bacterial inactivation.

Endotoxin-free flagellin was purified and depleted in endotoxin as described previously ^21^. To evaluate the protective effect, 5μg of flagellin was administrated intraperitoneally (i.p.) just before bacterial challenge. For IL-22 neutralizing experiment, mice received 200μg of neutralizing anti-IL-22 (AM22) or control isotype (a mouse IgG2a) antibodies intravenously 5 minutes before infection.

### Sample collection and processing

Mice were sacrificed 24h and 48h post-infection by NTHi. Broncho-alveolar Lavage (BAL) fluids, lungs, spleen and blood were collected and kept on ice till the processing or immediately frozen in liquid nitrogen.

BAL was performed by instilling 5 × 0.5 ml of sterile PBS + 2% fetal bovine serum (FBS) via a 1 ml sterile syringe with 23-gauge lavage needle into a tracheal incision. BAL samples were used for cytokine analysis, flow cytometry analysis and numbering of CFU. Lung tissues were collected aseptically and analyzed for CFU counts, cytokines, pulmonary cell profiles (flow cytometry analysis and lung cell restimulation) and histology. Blood was used for the determination of CFU counts.

### Flow cytometry

Cells harvested from BAL and lungs were washed and incubated with antibodies (BD, Franklin lakes, NJ, USA) for 30 min. Staining was performed as previously reported by Sharan et al. ^14^. Data were acquired on a LSR Fortessa (BD Biosciences) and analyzed with FlowJo™ software v7.6.5 (Stanford, CA, USA). Gating strategies are. Debris were excluded according to size (FSC) and granularity (SSC). Immune cells expressing CD45 were gated to analyse frequency, activation and number of cell subsets. Phenotypes are shown in the table 1.

**Table 1:**
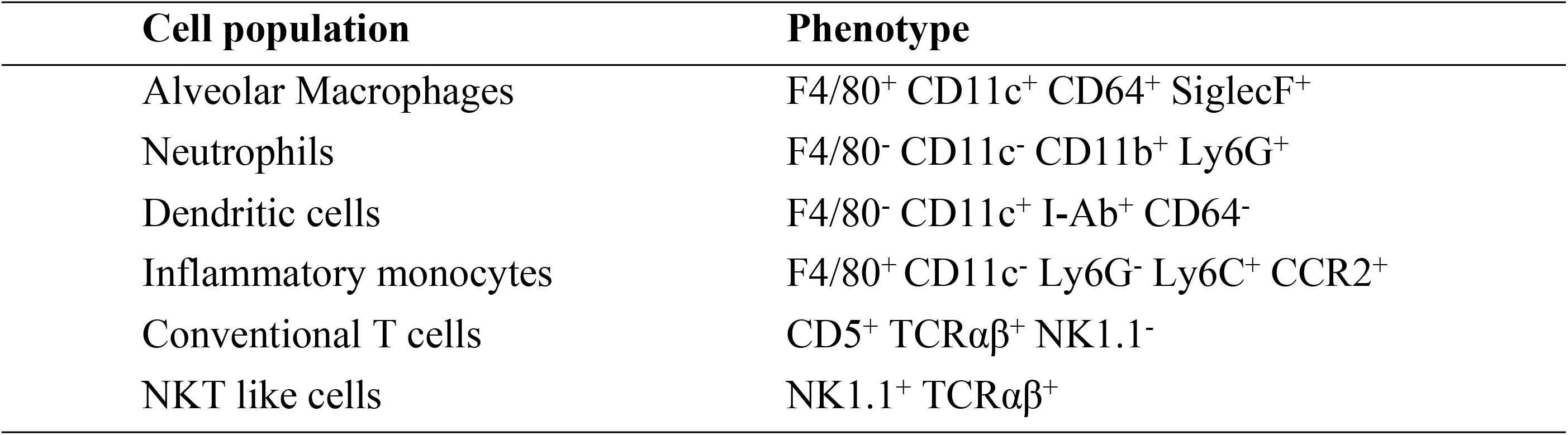
Phenotype of the major cell populations identified in this report.

### Cytokine measurement

Levels of IFN-γ, IL-1β, IL-6, IL-17, IL-22, IL-23 and tumor necrosis factor alpha (TNF-α) were quantified in lung tissue lysates and BAL using commercial ELISA kits (Invitrogen, San Diego, USA; Biotechne, Minneapolis, USA) (Table 2). Similarly, levels of IFN-γ, IL-17, and IL-22 were measured in the supernatants of dissociated lung cells (0.5×10^6^ of cells) re-stimulated with HK NTHi or not during 72h.

**Table 2:** List of the antibodies and of the ELISA kits used in this study.

| Flow cytometry mAb | Target | Manufacturer | Catalog Nb |
| --- | --- | --- | --- |
|  | FITC- I-Ab | Miltenyi Biotech | 130-102-168 |
|  | PE-F4/80 | Miltenyi Biotech | 130-102-422 |
|  | PerCP-Cy5.5 - CD103 | BD Biosciences | 563637 |

|  | PE-Cy7 - CD11c | BD Biosciences | 558079 |
| --- | --- | --- | --- |
|  | APC - CCR2 | Miltenyi Biotech | 130-119-658 |
|  | AF700 - CD86 | BD Biosciences | 560581 |
|  | APC-H7- Ly6G | BD Biosciences | 560600 |
|  | V450 - CD11b | BD Biosciences | 560455 |
|  | VioGreen - CD45 | Miltenyi Biotech | 130-110-665 |
|  | BV605 - Ly6C | Biolegend | 128036 |
|  | BV786 - CD64 | BD Biosciences | 741024 |
|  | PE-CF594 - SiglecF | BD Biosciences | 562757 |
|  | FITC - CD5 | Miltenyi Biotech | 130-102-574 |
|  | Tetramer mCD1d 167ms | NIH facility | 30663 |
|  | PerCP-Cy5.5 - NK1.1 | Miltenyi Biotech | 130-103-963 |
|  | PE-Cy7 - CD4 | Miltenyi Biotech | 130-102-411 |
|  | APC - CD25 | Miltenyi Biotech | 130-102-550 |
|  | AF700 - CD69 | BD Biosciences | 561238 |
| | APC-Vio770 - TCR$\gamma\delta$ | Miltenyi Biotech | 130-104-016 |
| | VioBlue -TCRβ | Miltenyi Biotech | 130-104-815 |
|  | V500 - CD8 | BD Biosciences | 130-109-252 |
|  | BV605 - CD45 | Biolegend | 103140 |
| ELISA kits | Target | Manufacturer | Catalog Nb |
| | IFN-γ ELISA kit | Invitrogen | 88-7314-88 |
| | IL-1β DuoSet | Biotechne | DY401 |
|  | IL-6 ELISA kit | Invitrogen | 88-7064-88 |
|  | IL-17 ELISA kit | Invitrogen | 88-7371-88 |
|  | IL-22 DuoSet | Biotechne | DY582 |
|  | IL-23 ELISA kit | Invitrogen | 88-7230-88 |
| | TNF-α ELISA kit | Invitrogen | 88-7371-88 |

### RT-PCR quantification of mRNA expression

Quantitative RT-PCR was performed to quantify mRNA of interest (Table 3). Results were expressed as mean ± SEM of the relative gene expression calculated for each experiment in folds (2^−ΔΔCt^) using *Gapdh* as a reference, and compared to non-infected PBS-treated control mice.

**Table 3:**
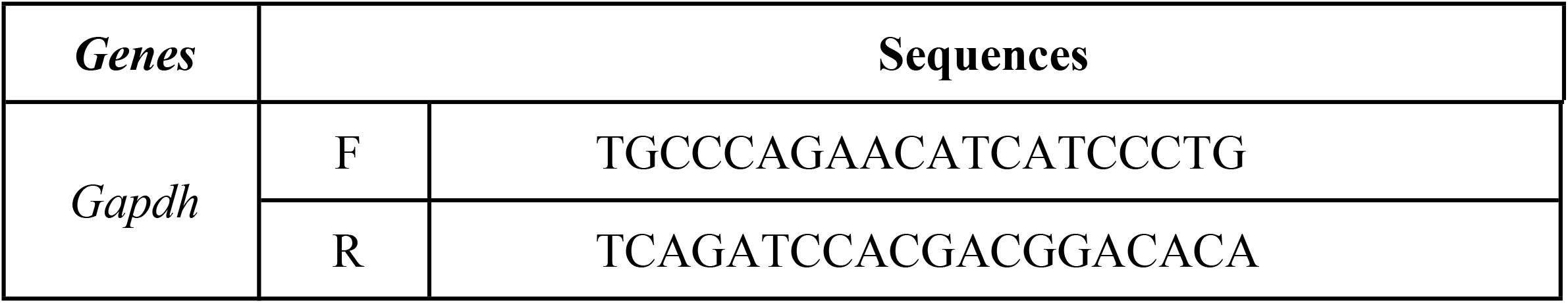
Primer sequences for qRT-PCR in mice. Forward (F) and reverse (R) primers are cited.

**Fig 1.**
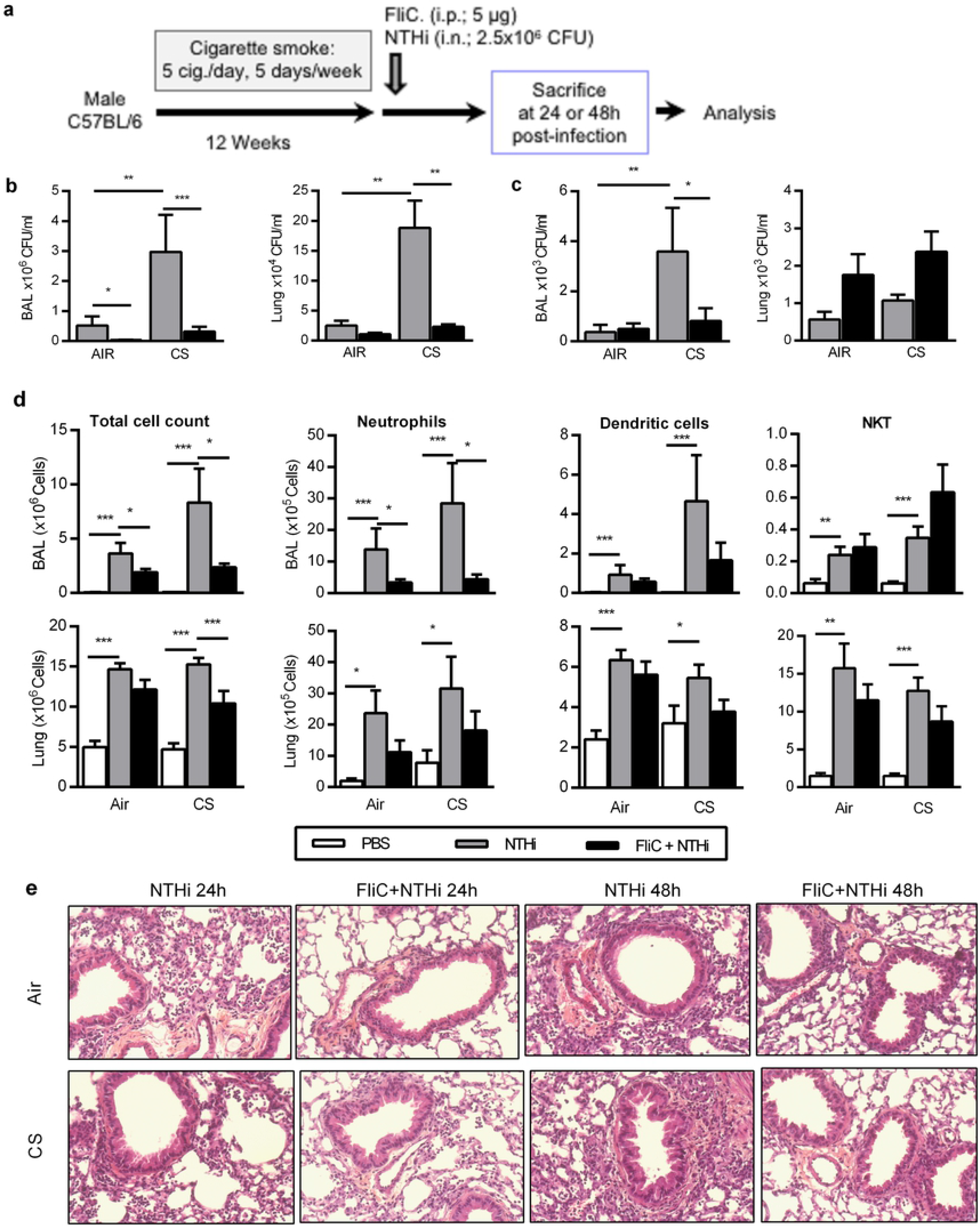

**Fig 2.**
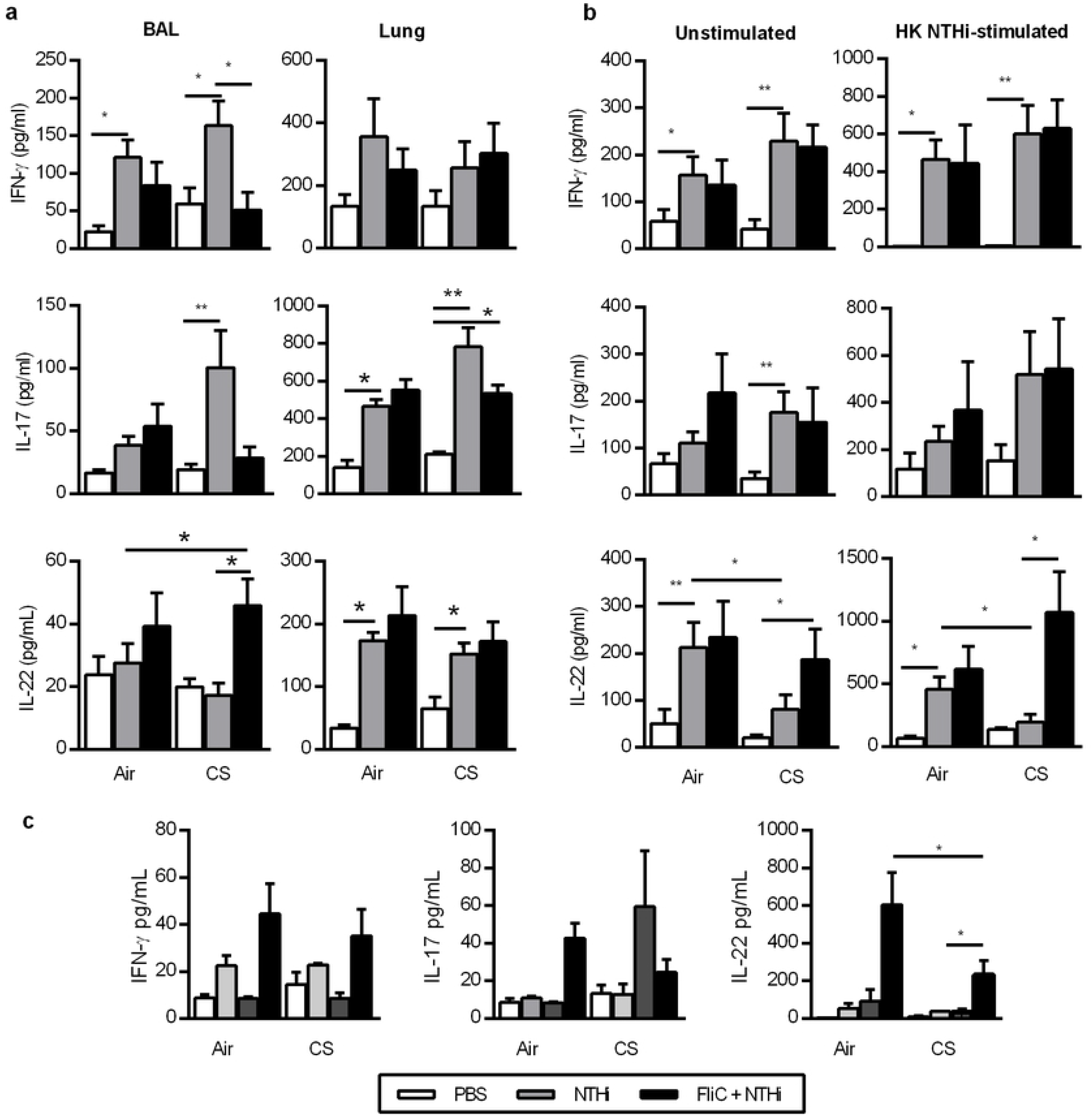

**Fig 3.**
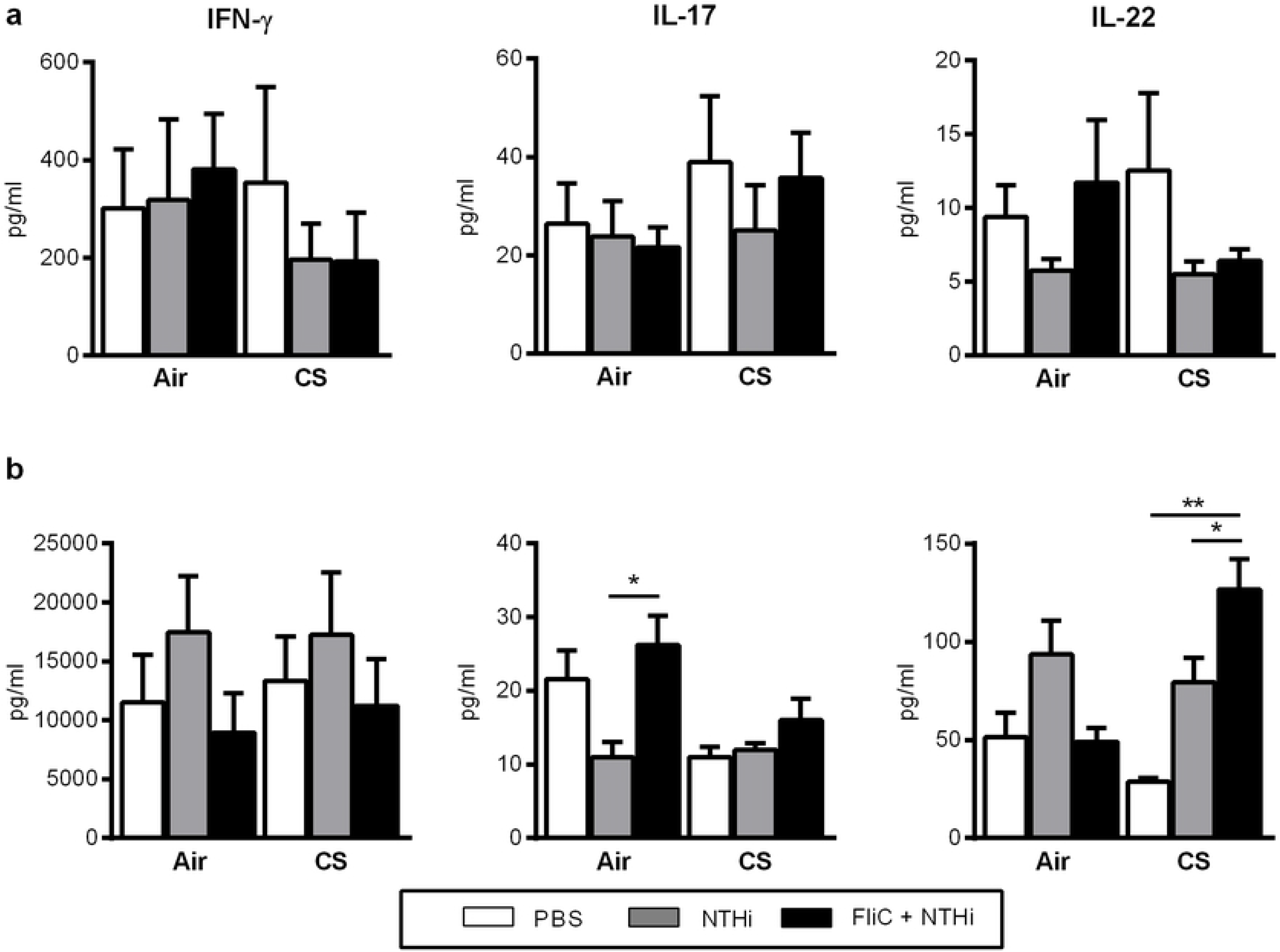

**Fig 4.**
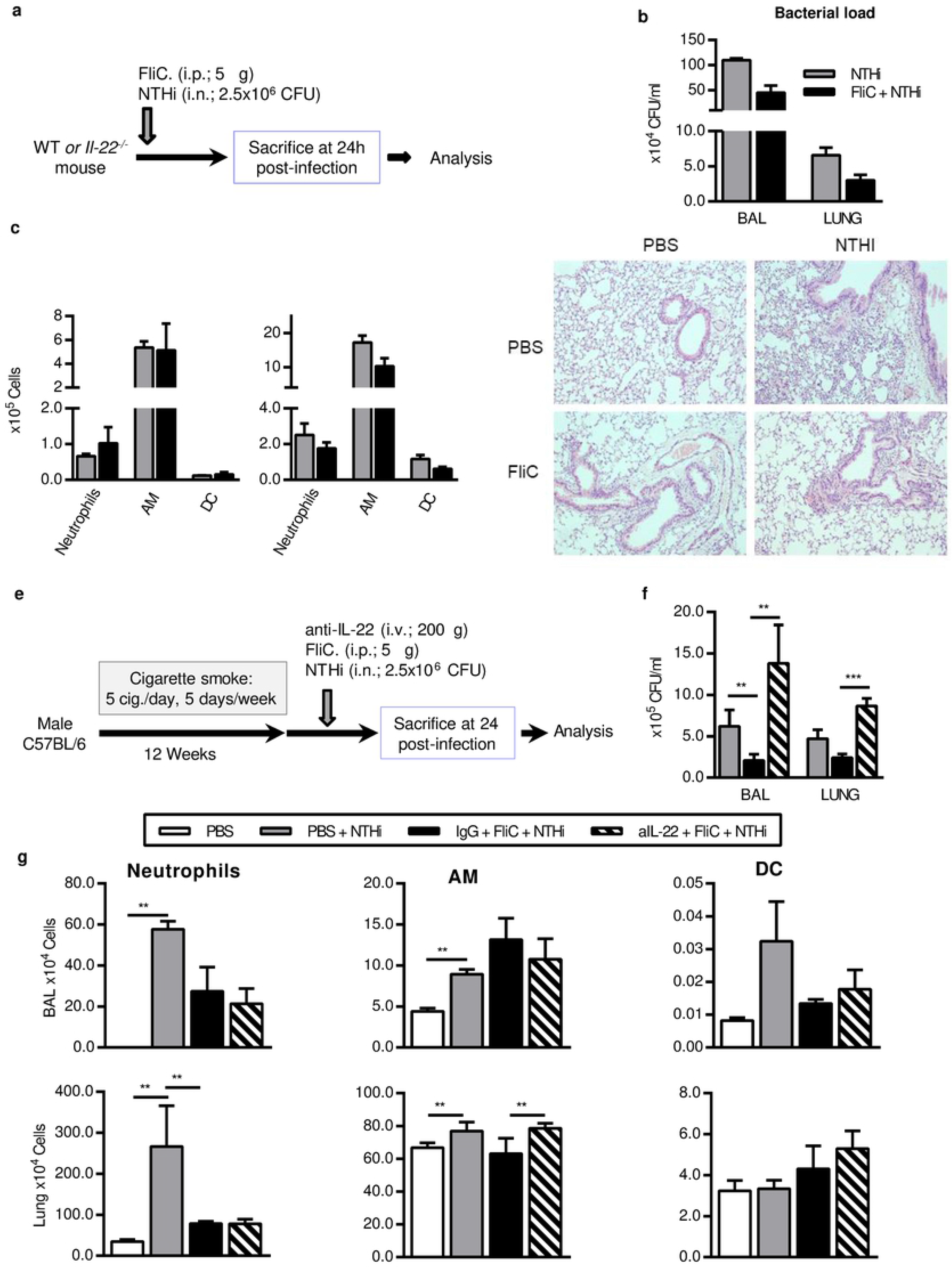

**Fig 5.**
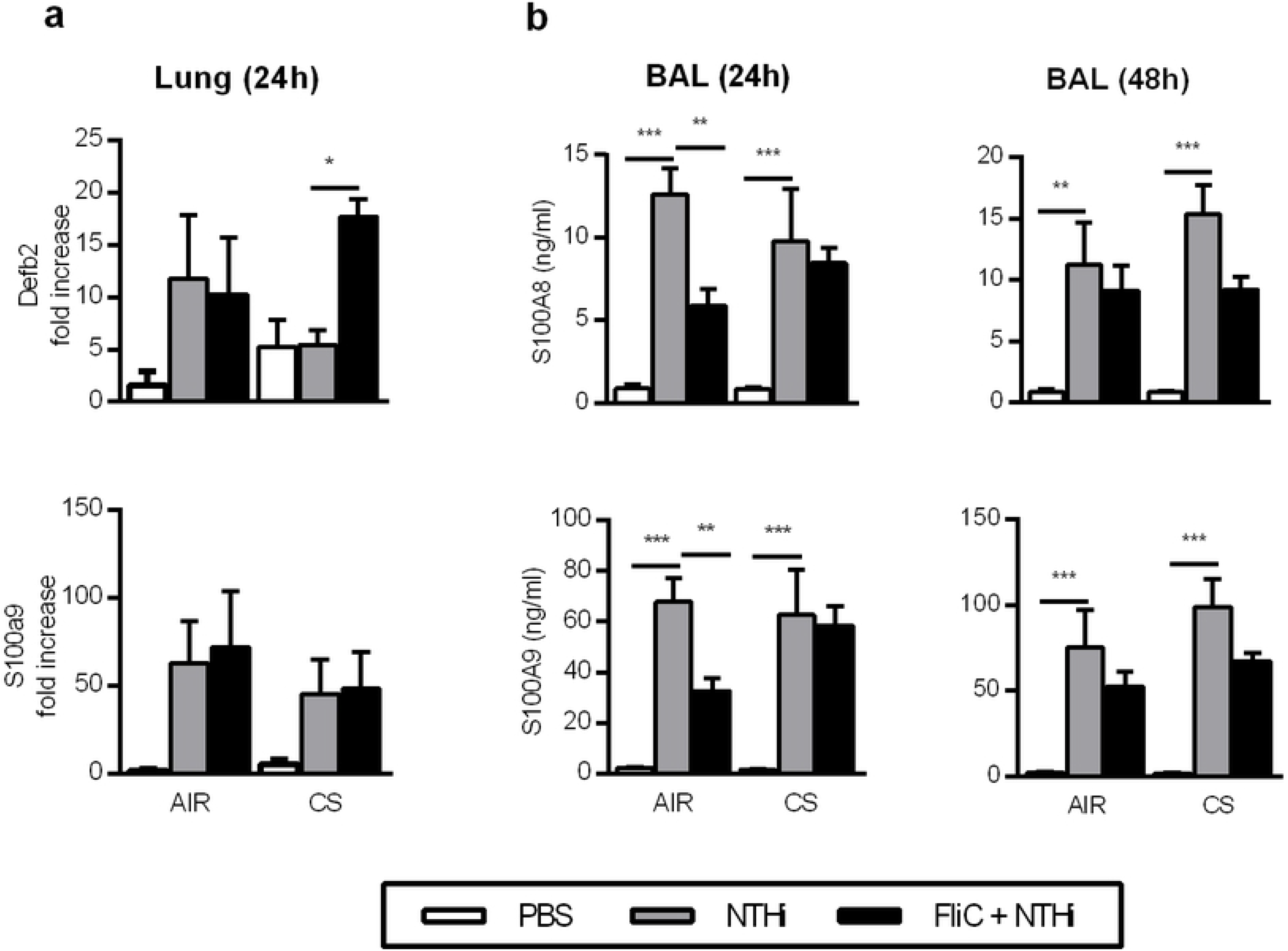

